# Deep homologies in chordate caudal central nervous systems

**DOI:** 10.1101/2024.06.03.597227

**Authors:** Matthew J. Kourakis, Kerrianne Ryan, Erin D. Newman-Smith, Ian A. Meinertzhagen, William C. Smith

## Abstract

Invertebrate chordates, such as the tunicate *Ciona*, can offer insight into the evolution of the chordate phylum. Anatomical features that are shared between invertebrate chordates and vertebrates may be taken as evidence of their presence in a common chordate ancestor. The central nervous systems of *Ciona* larvae and vertebrates share a similar anatomy despite the *Ciona* CNS having ∼180 neurons. However, the depth of conservation between the *Ciona* CNS and those in vertebrates is not resolved. The *Ciona* caudal CNS, while appearing spinal cord-like, has hitherto been thought to lack motor neurons, bringing into question its homology with the vertebrate spinal cord. We show here that the *Ciona* larval caudal CNS does, in fact, have functional motor neurons along its length, pointing to the presence of a spinal cord-like structure at the base of the chordates. We extend our analysis of shared CNS anatomy further to explore the *Ciona* “motor ganglion”, which has been proposed to be a homolog of the vertebrate hindbrain, spinal cord, or both. We find that a cluster of neurons in the dorsal motor ganglion shares anatomical location, developmental pathway, neural circuit architecture, and gene expression with the vertebrate cerebellum. However, functionally, the *Ciona* cluster appears to have more in common with vertebrate cerebellum-like structures, insofar as it receives and processes direct sensory input. These findings are consistent with earlier speculation that the cerebellum evolved from a cerebellum-like structure, and suggest that the latter structure was present in the dorsal hindbrain of a common chordate ancestor.

## Introduction

The human nervous system is the most complex biological structure we know in the universe. To gain a better understanding of the origin of its complexity, researchers turn to organisms with less complex nervous systems. Yet, even among the simplest vertebrates, the cyclostomes (*e.g.*, lamprey and hagfish), we find central nervous systems (CNS) that already have the core organization, and most of the neuron types, found in mammalian CNSs (*1*, *2*). Invertebrate chordates, tunicates and cephalochordates, may allow us to explore still deeper origins of the neuron types and structures of the vertebrate CNS. Tunicates, the sister group to the vertebrates (*3*), have proven to be particularly tractable experimental biological models, with the solitary ascidian *Ciona* being particularly well studied. The CNS of a *Ciona* larva follows a similar developmental program to that of a generalized vertebrate CNS, and once formed has distinct anterior and posterior domains mirroring the vertebrate forebrain, midbrain, midbrain-hindbrain junction (MHJ), hindbrain and spinal cord [Table 1A; (*4*)]. Historically, these brain regions in tunicate larvae (including *Ciona*) were called the *anterior sensory vesicle*, *posterior sensory vesicle*, *neck*, *motor ganglion* and *caudal nerve cord*, respectively. While it was once thought that tunicate larvae lacked distinct domains with homology to vertebrate fore- and midbrain regions, recent functional and gene expression studies, aided by findings from the *Ciona* connectome (*5*), have resolved the issue with the anterior sensory vesicle showing homology to the vertebrate forebrain, and the posterior sensory vesicle showing homology to the midbrain. For example, the anterior sensory vesicle mirrors the developing vertebrate forebrain in expressing the genes *Otx* and *Dmrt1*, and has a dopamine-expressing domain that appears to be homologous to the hypothalamus (*6–10*). By contrast, the posterior sensory vesicle receives, integrates, and relays sensory inputs, mirroring the vertebrate midbrain (*5*, *11*, *12*). However, homology between the tunicate motor ganglion and caudal nerve cord and the vertebrate hindbrain and spinal cord is in both cases still in question. Specifically, the apparent absence of motor neurons in the caudal nerve cord, as well as the expression of several genes in the motor ganglion that are expressed in the vertebrate spinal cord, led to speculation that the motor ganglion had properties linking it to both the vertebrate hindbrain and spinal cord (*13–16*). We present here results showing that the *Ciona* caudal nerve cord has previously unrecognized motor neurons along its length, consistent with spinal cord homology. In addition, we present evidence that a cluster of neurons in the dorsal motor ganglion have properties pointing to their shared origin with the vertebrate cerebellum, supporting homology between the motor ganglion and vertebrate hindbrain. In a recent publication (*17*) we substituted tunicate-specific larval brain nomenclature in favor of their proposed vertebrate homologs (*e.g.*, “hindbrain” instead of “motor ganglion”). This was done to reflect the growing data supporting homology between these brain regions, and to help make the text more easily understandable to a broader readership.

**Table 1.**
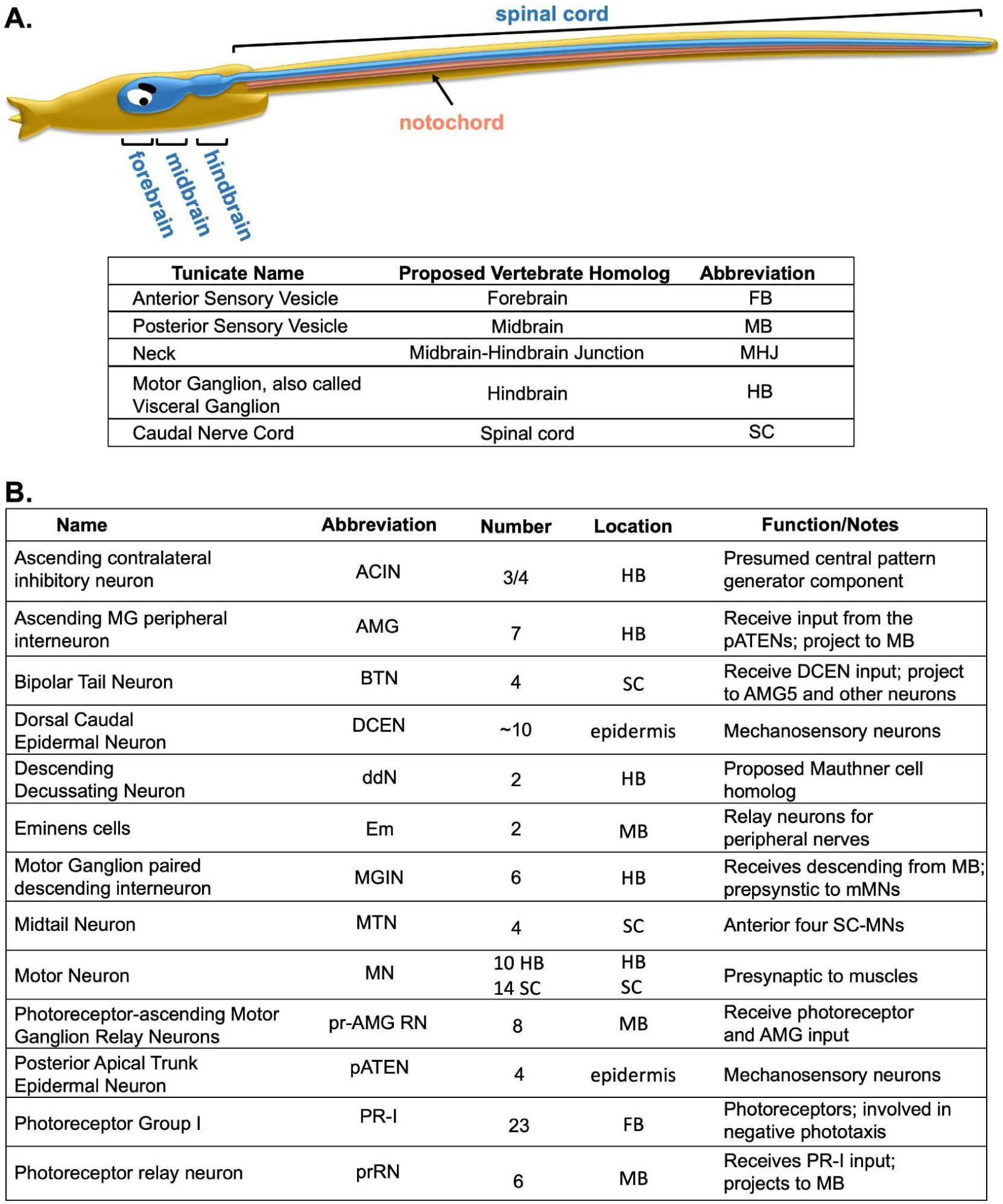
A. Proposed tunicate-to-vertebrate homology of CNS domains. B. Summary of neuron types mentioned in the text.

However, because one of the primary aims of this manuscript is to resolve ambiguity around the nature of the tunicate motor ganglion and caudal nerve cord, we will retain the tunicate names but conclude that the use of the vertebrate-counterpart names “hindbrain” and “spinal cord” is now justified, and extends congruence between the vertebrate and tunicate nervous systems.

Table 1A is provided to assist the reader to translate between the tunicate and vertebrate nomenclatures.

## Results

### Left/right paired *Mnx* and *VACHT* co-expressing cells are present along the length of the Ciona *larval caudal nerve cord*

Using *in situ* hybridization in *Ciona* larvae we observed twelve cells in the caudal nerve cord expressing the cholinergic marker VACHT (Figure 1A, yellow asterisks). These cells are distributed along the length of the tail and lie posterior to the caudal-most motor neurons of the motor ganglion (MN5L and R). We find that these neurons co-express the motor neuron-specific transcription factor *mnx* (Figure 1B-C), suggesting homology to vertebrate motor neurons.

**Figure 1.**
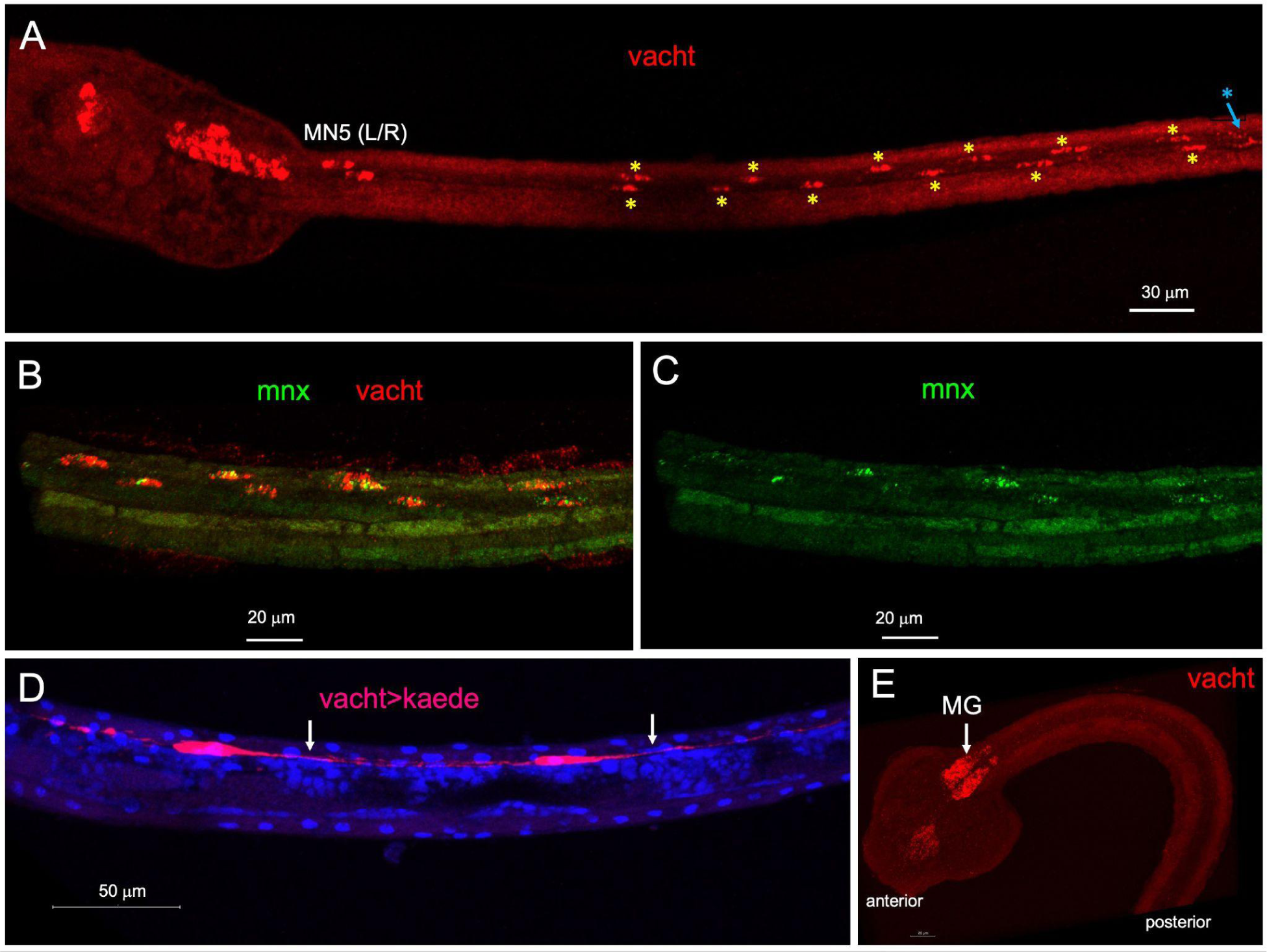
Motor neurons in the caudal nerve cord. **A**. *In situ* hybridization for VACHT in a *Ciona* larva. Yellow asterisks indicate the twelve tail motor neurons. The most posterior pair of these [MN5 Left (L) and Right (R)] are indicated. Blue asterisk indicates a bipolar tail neuron. **B**. Tail of a *Ciona* larva showing overlay of *in situ* hybridization for VACHT and mnx. **C**. Single channel for mnx from panel B. **D**. Caudal nerve cord motor neurons expressing kaede fluorescent protein under control of the VACHT promotor. Arrows indicate projections from the motor neurons. Shown is the result of staining with anti-kaede antibodies and Dapi. **E**. In situ hybridization for VACHT to tailbud stage embryo. MG = motor ganglion.

Expression of the fluorescent protein Kaede from the VACHT promoter allows us to visualize the morphology of the tail motor neurons with their posteriorly-extending axons (arrows, Figure 1D). A similar pattern was reported previously in *Ciona* transgenics expressing GFP under the control of the VACHT promoter, but was not further characterized (*18*, *19*). Despite this, it is widely reported that the *Ciona* caudal nerve cord either lacks motor neurons, or is completely devoid of neurons altogether, having only ependymal cells (*4*, *14*, *18*, *20*, *21*). This appears to be explained, at least in part, by the relatively late development of the motor neurons in the caudal nerve cord. For example, at the mid-tailbud stage, a common temporal endpoint for expression studies, VACHT expression detected by *in situ* hybridization is observed in the motor ganglion (MG; Figure 1E), but not yet in the caudal nerve cord.

### Caudal nerve cord *Mnx* and *VACHT* co-expressing cells evoke muscle contraction

In order to assess the function of caudal nerve cord motor neurons, embryos were electroporated (*22*) with the red-shifted channelrhodopsin *chrimson* under the control of the VACHT promoter (*VACHT>Chrimson-mCherry*). Control samples did not receive the transgene. The resulting electroporated and control embryos were then bisected at the early-tailbud stage [stage 20, (*23*)] between the tail and trunk in order to isolate the developing caudal nerve cord motor neurons from the motor ganglion motor neurons. In addition, because axonal projections are not observed from the motor ganglion motor neurons until the mid-tailbud stage (*24*), the resulting tail fragments were predicted not to have axonal projections originating from the motor ganglion. The tail fragments were then cultured to the larval stage (stage 26; approximately eight hours after bisection) at which point they were assessed. For the assay, 30-second movies of the electroporated and control larval-stage tail fragments were recorded at 8.9 frames/second with continuous 490 nm LED illumination (*i.e.*, outside the activation range of Chrimson). At the midpoint of the recording, a 590 nm LED lamp was turned on to activate the Chrimson. Figure 2 shows five-second temporal projections of the movies, immediately before (Figure 2A) or immediately after (Figure 2B) the 590 nm illumination (*n* = 7 for *VACHT>Chrimson-mCherry* expressing tail fragments; and n=9 for control tail fragments). Six of the seven *VACHT>Chrimson-mCherry* expressing tail fragments responded by beating immediately following illumination with the 590 nm lamp, while none of nine control tail fragments responded (Figure 2 and Video 1). Three repetitions of the assay were performed with the tail fragments, with identical results (data not shown). At the completion of the assay, the tail fragments were fixed and immunolabeled with anti-mCherry antibodies. A representative tail fragment is shown in Figure 2C with the caudal nerve cord motor neurons marked with red arrows. SFigure 1 shows the immunolabeling results for six of seven expressing tail fragments (one having been lost after the assay). Based on these results, we conclude that mnx/vacht-positive cells in the caudal nerve cord are functional motor neurons.

**Figure 2.**
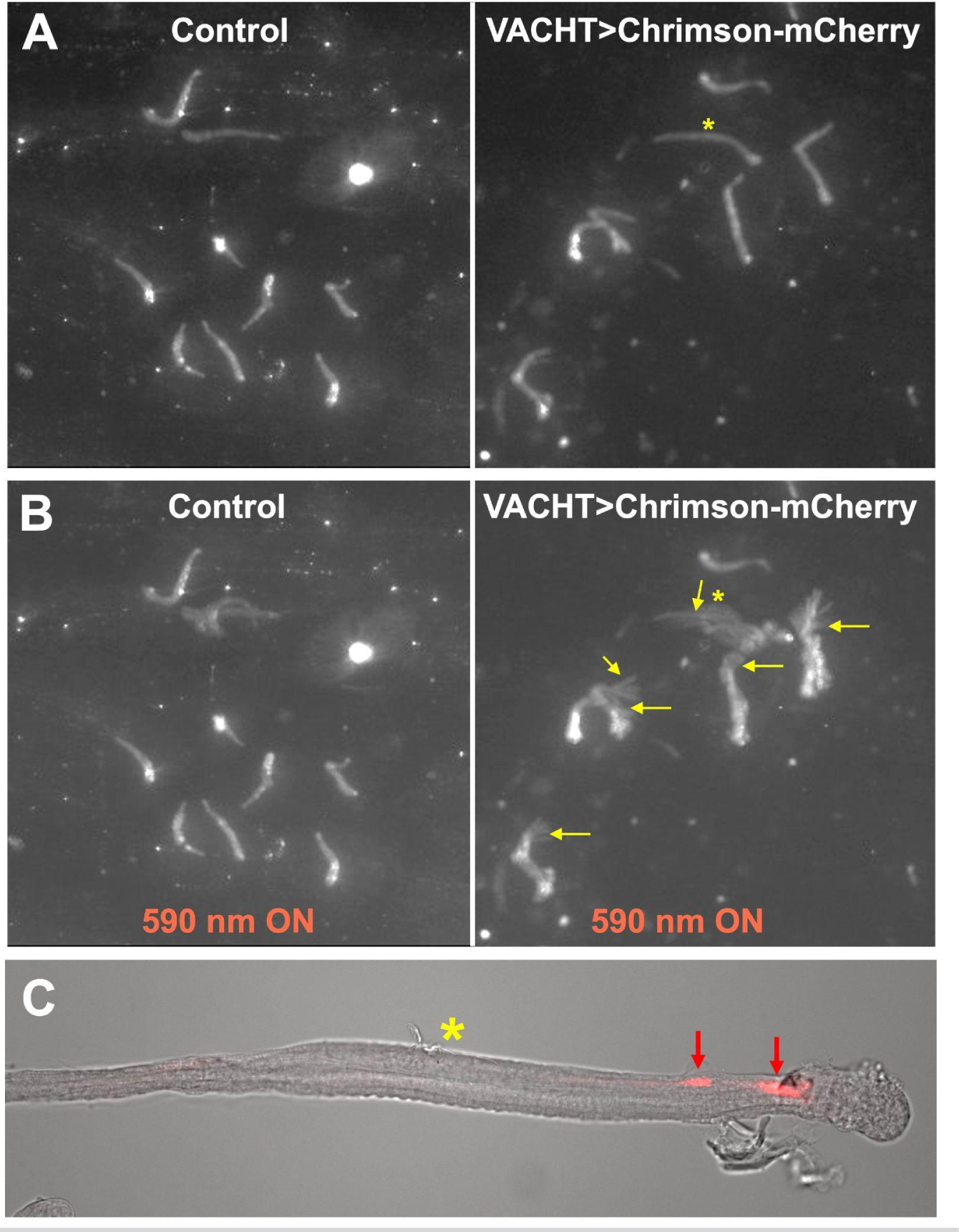
Functional assessment of caudal nerve cord motor neurons. **A** and B. Five-second time projections of tail fragments immediately before (A) and after (B) a 590 nm LED lamp was illuminated to activate chrimson. Yellow arrows indicate tail beating. C. Example of VACHT>Chrimson-mCherry expressing tail fragment immunolabeled with anti-mCherry antibody. Yellow asterisk indicates the corresponding tail fragment in panels A and B, while red arrows indicate caudal nerve cord motor neurons. Anterior is to the right.

### Caudal nerve cord motor neurons and the *Ciona* connectome

The analysis of neurons in the *Ciona* connectome project did not extend caudally past the approximate midpoint of the caudal nerve cord (*5*). Nevertheless, the analysis did include four caudal nerve cord neurons named *midtail neurons* (MTNs) (bottom panel, Figure 3A). The MTNs were tentatively identified as motor neurons, although they have not been described in detail [see *Supplemental data, Fig. 1-data1-v1* in (*5*)]. A comparison between our VACHT *in situ* hybridization results (top panel, Figure 3A) and a reconstructed larvae from the connectome serial-section electron micrographs (bottom pane, Figure 3A) reveals the MTNs (bottom) and the rostral-most VACHT-expressing caudal motor neurons (top) occupy similar positions in the CNS. This is particularly evident in the spacing between MN5 of the motor ganglion and the MTNs/caudal nerve cord motor neurons in two samples (GAP, Figure 3A). In addition, the morphology of the MTNs, with short irregularly-extending axons, matches the morphology of caudal nerve cord motor neurons shown in Figure 1D. We conclude that the MTNs and the caudal nerve cord motor neurons identified in Figure 1A are the same. Details of the four MTNs from the EM reconstructions shows that they send short descending axons within the caudal nerve cord and are presynaptic to muscle cells (Figure 3B-G), as well as to the basement membrane opposite the notochord (Figure 3F), and to other neurons with axons in the tail, including ddNs, and MN2 (SFigure 2).

**Figure 3.**
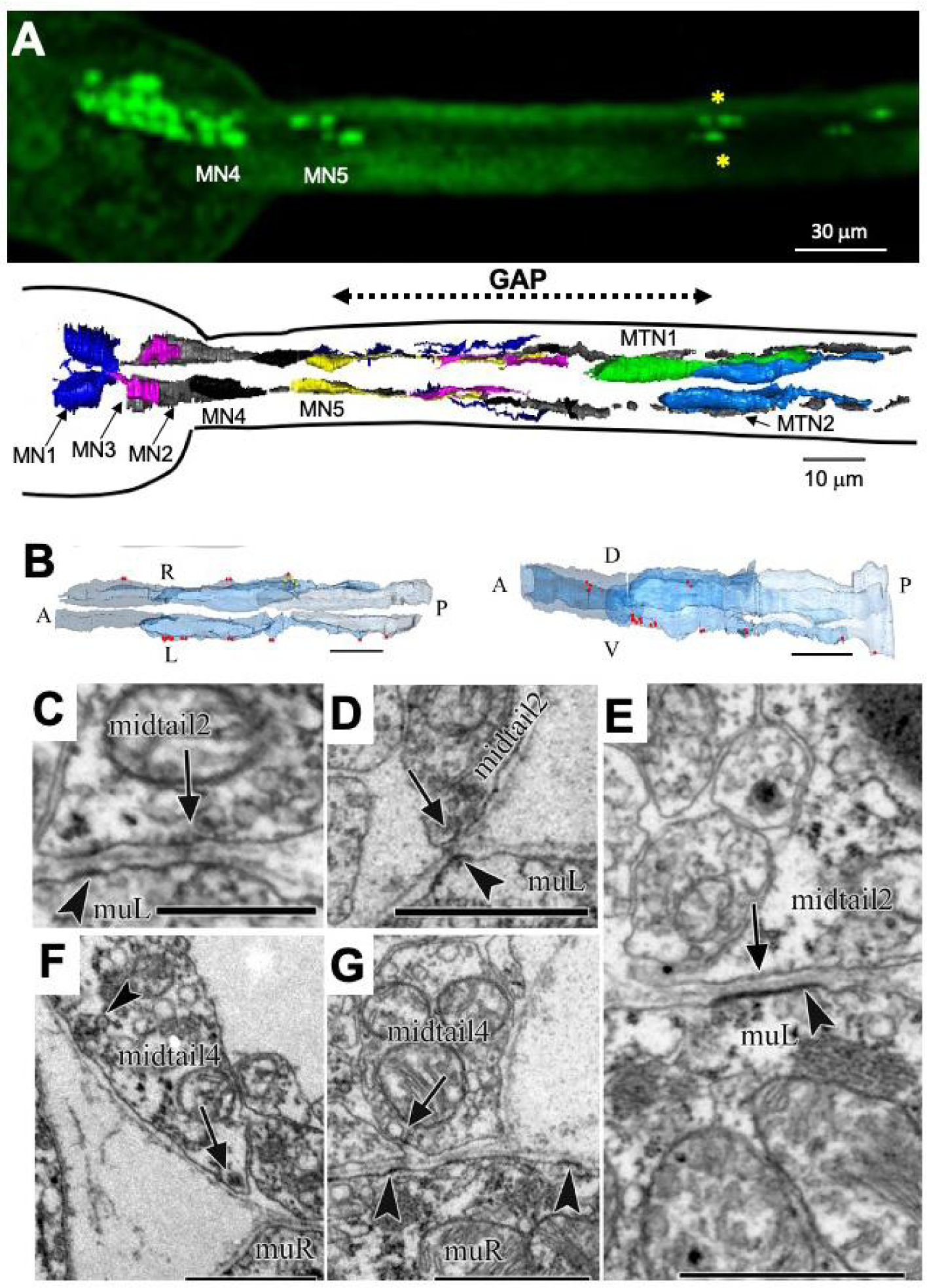
Midtail motor neurons. **A**. Comparison between the anterior/posterior positions of motor ganglion motor neurons (MN4 and MN5) and the caudal nerve cord motor neurons (yellow asterisks), as detected by *in situ* hybridization (top), and between the positions of MN4 and 5 and the midtail motor neurons (MTN1 and MTN2) from the connectome (bottom). **B**. Dorsal and left lateral views of midtail neurons with neuromuscular junctions marked by red puncta; yellow puncta represent synapses onto the basement membrane. **C-E.** Putative neuromuscular synapses (arrows) from MTN2 onto left dorsal muscle cells (muL). **F-G**. Putative neuromuscular synapses (arrows) from MTN4 onto right dorsal muscle cells (muR) Postsynaptic densities are present on muscle cells (arrowheads in C-E; G). Some presynaptic sites have putative vesicles of mixed sizes (C, D and F). Synapse onto basement membrane shown by arrowhead in F. Scale bars: 10μm (B); 1μm (C-G). (muL, muR = muscle Left and Right).

Given that the caudal nerve cord motor neurons and the MTNs identified through the connectome data are determined to be the same, we can use the connectome data to analyze their connectivity. The MTNs are postsynaptic to many of the same interneurons as are motor neurons in the motor ganglion, and they appear to be components of previously characterized sensorimotor circuits. For example, Figure 4A shows the core neural circuit for negative phototaxis with neurons grouped according to type (*5*, *25*). In this circuit, the twenty-three Group-I photoreceptors (PR-I) project primarily to two classes of relay interneurons in the midbrain (six prRNs and eight pr-AMG RNs). These midbrain relay neurons in turn project to the left and right *motor ganglion interneurons* (MGINs; three on each side). As previously reported, the MGINs are then presynaptic to the ten motor neurons of the motor ganglion. However, the connectome shows that MGINs are presynaptic to the MTNs as well (Figure 4b). At the level of individual neurons, it is clear that the left-sided MTN2 is postsynaptic to MGIN-2L and MN2L, while both left MTNs (2 and 7) are postsynaptic to the right ddN (Figure 4B). The ddNs have been compared to the Mauthner cells of fish and amphibians, which mediate the startle response (*26*). By contrast, the two right MTNs (1 and 4) are postsynaptic to both MGIN-2R, MN2R, and the left ddN (Figure 4B). Significantly, of the ten motor ganglion motor neurons, the MTNs are targeted only by the left/right MN2s. This pair of motor neurons shows spontaneous and rhythmic spiking activity, and is thought to be a central component of the *Ciona* larval swim central pattern generator (CPG) (*24*, *27*). In fact, MTN2 and MTN7 make reciprocal synapses with the two MN2s (red asterisks Figure 4b). Additional components of the CPG are thought to be the *ascending contralateral inhibitory neurons* (ACINs) (*18*, *28*). These are thought to act primarily at the level of the MGINs (*5*), and are predicted by the connectome to target the left and right MGIN-2s, which are presynaptic to the MTNs (Figure 4B). However, the connectome provides information on synaptic connectivity only for the rostral-most four of the 12 caudal nerve cord motor neurons, and, thus, it remains to be determined whether the remaining motor neurons have similar connectivity.

**Figure 4.**
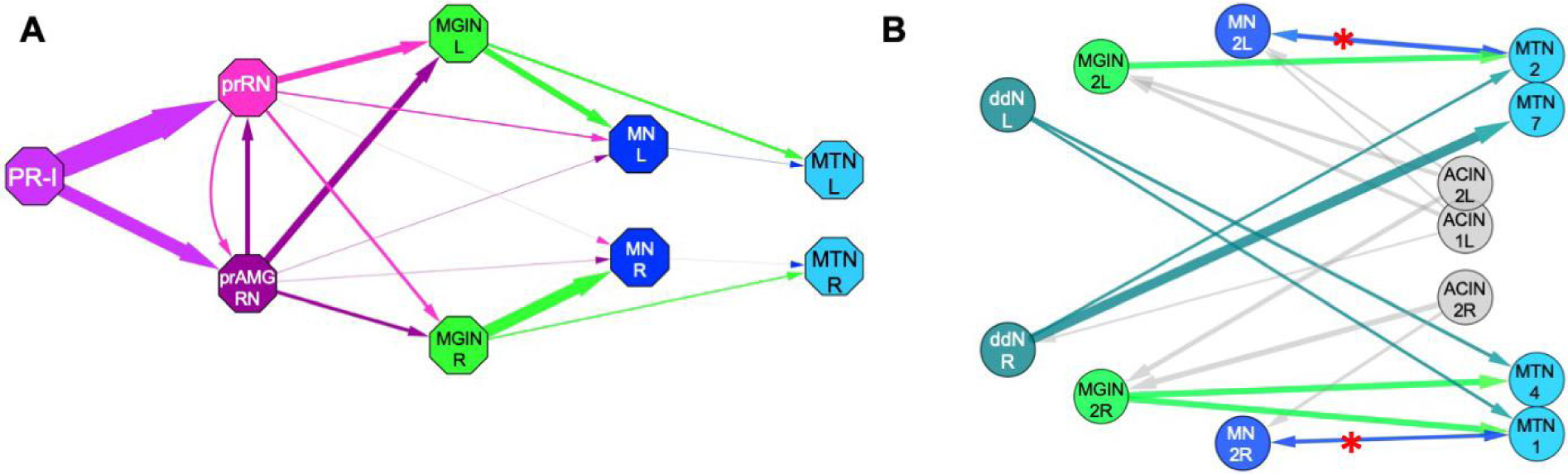
Synaptic connections of hindbrain and caudal (midtail) motor neurons. **A**. Example of a *Ciona* sensorimotor circuit (for negative phototaxis) emphasizing descending projections to both motor ganglion motor neurons and the MTNs. In the diagram, neurons of each class (*e.g.*, twenty-three photoreceptors) are grouped (octagons) and their synaptic contact strengths are summed and represented by the widths of the arrows. **B**. Synaptic inputs to the four caudal motor neurons. Circles represent individual neurons. Red asterisks indicate reciprocal synapses. **Abbreviations**: PR = photoreceptor; prRN = photoreceptor relay neuron, prAMG RN = photoreceptor-ascending motor ganglion relay neuron; MGIN = motor ganglion interneuron; MN = motor ganglion motor neuron; MTN = midtail neuron; ddN = descending decussating neuron; ACIN = ascending contralateral inhibitory neuron. All connectivity data, and the color scheme for neuron classes, is from (*5*).

#### The Ciona motor ganglion

As detailed in the Introduction, the *Ciona* motor ganglion has been equated with both the vertebrate hindbrain and the spinal cord. This is largely based on the presumed absence of motor neurons in the caudal nerve cord. Our description of motor neurons running the length of the caudal nerve cord (see above), along with its spinal cord-like morphology and development, argues that this structure in *Ciona*, and therefore presumably all tunicates, does indeed share true anatomical homology with the vertebrate spinal cord. Accordingly, should the motor ganglion be regarded as a strict homologue of the hindbrain? The motor ganglion contains five left-right pairs of motor neurons that express the genes *Phox2* and *Tbx20*, pointing to a common origin with vertebrate cranial motor neurons, rather than with spinal motor neurons (*14*). Moreover, as will be described below, a cluster of seven neurons, the AMGs in the dorsal motor ganglion, have apparent homology to a major vertebrate hindbrain domain - the cerebellum - furthering the link between the motor ganglion and the hindbrain.

The *Ciona* motor ganglion has distinct dorsal and ventral domains, with the ventral domain containing the above-mentioned motor neurons, MGINs, ddNs, and the ACINs, all of which mediate sensorimotor responses [see Table 1 and (*29*)]. By contrast, the dorsal domain of the motor ganglion is occupied exclusively by the seven AMG neurons (Figure 5A). Unlike the motor functions of the ventral motor ganglion (VMG), the AMGs process and relay input from the mechanosensory pATENs in the dorsal trunk epidermis to the midbrain (*30*). The AMG group consists of six inhibitory/VGAT-positive interneurons with a central excitatory/VACHT-positive neuron (AMG5) [Figure 5B and (*25*)]. A reconstruction of the motor ganglion from the serial sections of the connectome project highlights the anatomical separation of the AMG group from the ventral motor ganglion (VMG) (Figure 5C). Also shown in Figure 5C are the pATENs, as well as the primary targets of the AMGs in the midbrain (eminens neurons and pr-AMG RNs). The functional distinction between the AMG group and the VMG is particularly evident when their connectivities to each other, and to the midbrain, are compared (Figure 5D). While the VMG receives extensive descending input from the midbrain, the AMG group receives very little such input, and, rather, is characterized by its extensive ascending projections to the midbrain. Figure 5E shows the ascending projections from the AMG, using the *hox 10* cis-regulatory region to express GFP in AMGs 3, 4 and 5. While there is extensive synaptic connectivity within the AMG group and the VMG neurons (gray arrows in Figure 5D), there is very little between the two, and the few synaptic connections that are present consist almost exclusively of projections from the AMG group to the VMG.

**Figure 5.**
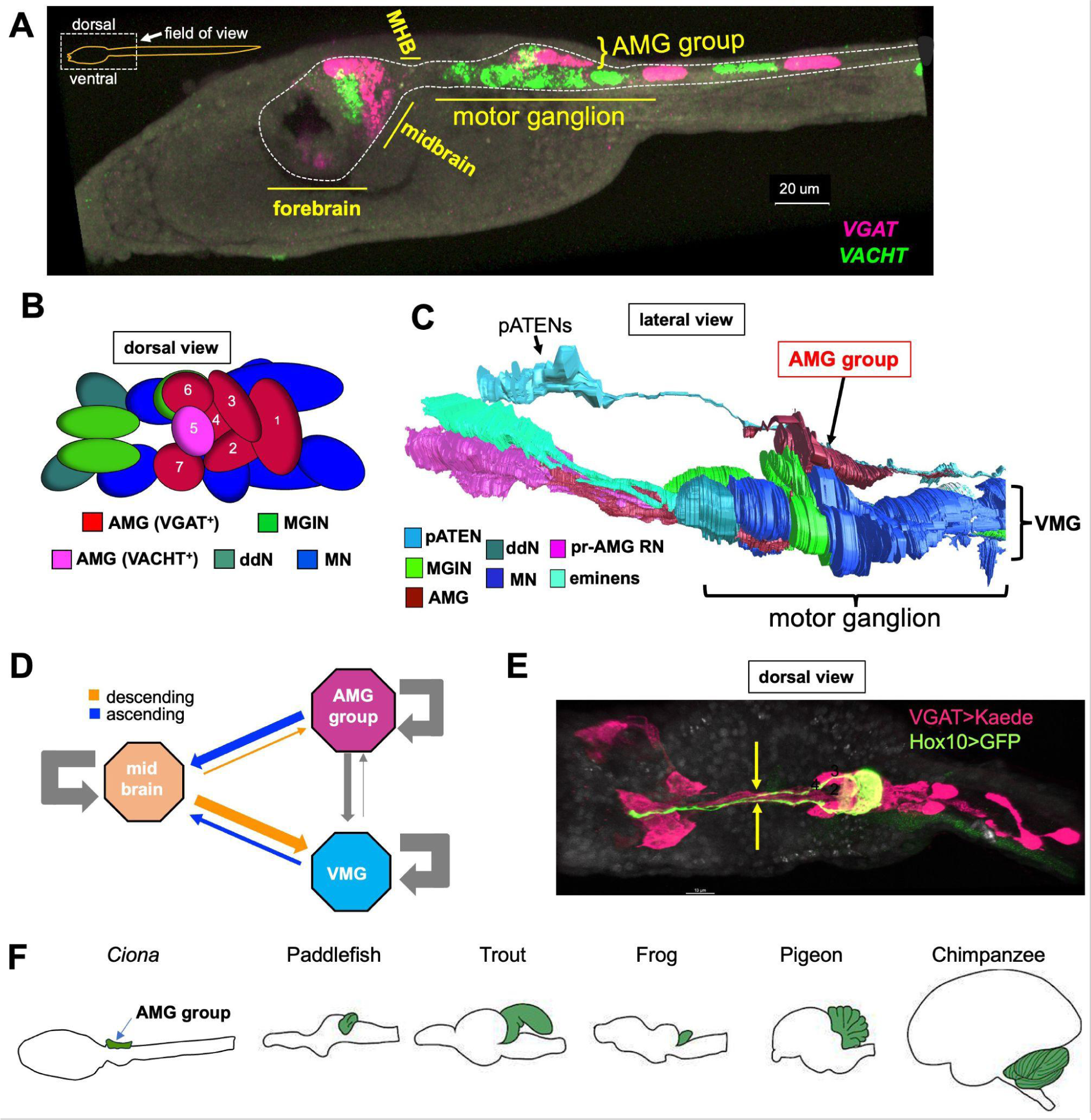
The *Ciona* AMG group. **A.** The *Ciona* larval nervous system (outlined). Image shows fluorescent *in situ* hybridization for VACHT and VGAT. The major CNS subdivisions, including the AMG group are indicated. **B**. Diagram of the motor ganglion (dorsal view). Neurons are shown as ovals that correspond to their sizes and relative positions from the connectome. **C.** Lateral view of the motor ganglion reconstructed from serial-section electron micrograph. Also shown are primary inputs to the AMGs, the *posterior apical trunk epidermal neurons* (pATENs), as well as the targets of the AMGs, the *photoreceptor-ascending motor ganglion neuron relay neurons* (pr-AMG RNs) and eminens cells. VMG= ventral motor ganglion. **D.** Summary of synaptic connectivity between the AMG group, the ventral motor ganglion (VMG) and the midbrain. The strengths of all synaptic connections between neurons in the brain regions were summed and then normalized by the number of neurons in each region. These values are reflected in the width of the arrows between the brain regions. **E**. Ascending projections from AMGs 2, 3, and 4 labeled with a Hox10 promoter>GFP plasmid (green). Also shown are VGAT-positive neurons (magenta). **F**. Relative position of the AMG group in the CNS (arrow) compared with those of cerebella (green) from various vertebrates. Image modified from (*31*).

The distinct features of the dorsal motor ganglion bring into question its evolutionary origin, and its relationship, if any, to brain regions of vertebrate nervous systems. Interestingly, the anatomical location of the AMG group resembles that of the vertebrate cerebellum (Figure 5F). Moreover, like the AMG group, the vertebrate cerebellum is distinct from the rest of the hindbrain both in its function and synaptic connectivity. In the following sections we will explore this relationship further and show a similar inductive pathway, circuit architecture and gene expression between the AMG group and the vertebrate cerebellum and cerebellum-like structures.

### Conserved inductive pathway for the AMG group and cerebellum

In vertebrates the cerebellum is induced via the action of FGF8 produced at the *midbrain-hindbrain boundary* (MHB) (*31*). The *Ciona* ortholog of this gene, called FGF8/17/18, shows a remarkably well-conserved expression pattern in the developing CNS, being expressed in the CNS neck region, immediately posterior to neurons expressing *OTX* and *engrailed (En)*, as in vertebrates (*32*). The naming of FGF8/17/18 reflects the multiple vertebrate orthologs of the single *Ciona* gene resulting from independent whole-genome duplications in the vertebrate lineage that followed the split from tunicate ancestors (*33*). Based on morphology, gene expression, and inductive activity the *Ciona* larval CNS neck region has been equated with the MHB (*8*, *15*). Previous studies showed that the loss of FGF8/17/18 disrupts motor ganglion gene expression (*34*), although disruptions specific to the AMG group were not addressed. To investigate a possible role for FGF in the development of the AMG group, *Ciona* embryos were treated with the FGF receptor inhibitor SU5402. To assess the presence/absence of the AMG group, experiments were performed in a stable transgenic line expressing Kaede fluorescent protein under the control of the VGAT promoter (*35*), which labels the six inhibitory AMG neurons (1-4, and 6-7). We found that treatment of the embryos for 30 minutes with 20 μmole SU5402 at the late neurula stage eliminated VGAT expression in the dorsal motor ganglion, while vehicle control (DMSO) had no effect (Figure 6A and B). Despite the loss of the AMGs, other VGAT-expressing CNS neurons appeared to be present, including those in the midbrain (red asterisks in Figure 6A and B) and the ACINs of the motor ganglion (yellow arrows), indicating that the AMGs are uniquely induced at this developmental stage. Treatment of older embryos with SU5402, from the initial tailbud stage onward, had no effect on the development of the AMG group (not shown).

**Figure 6.**
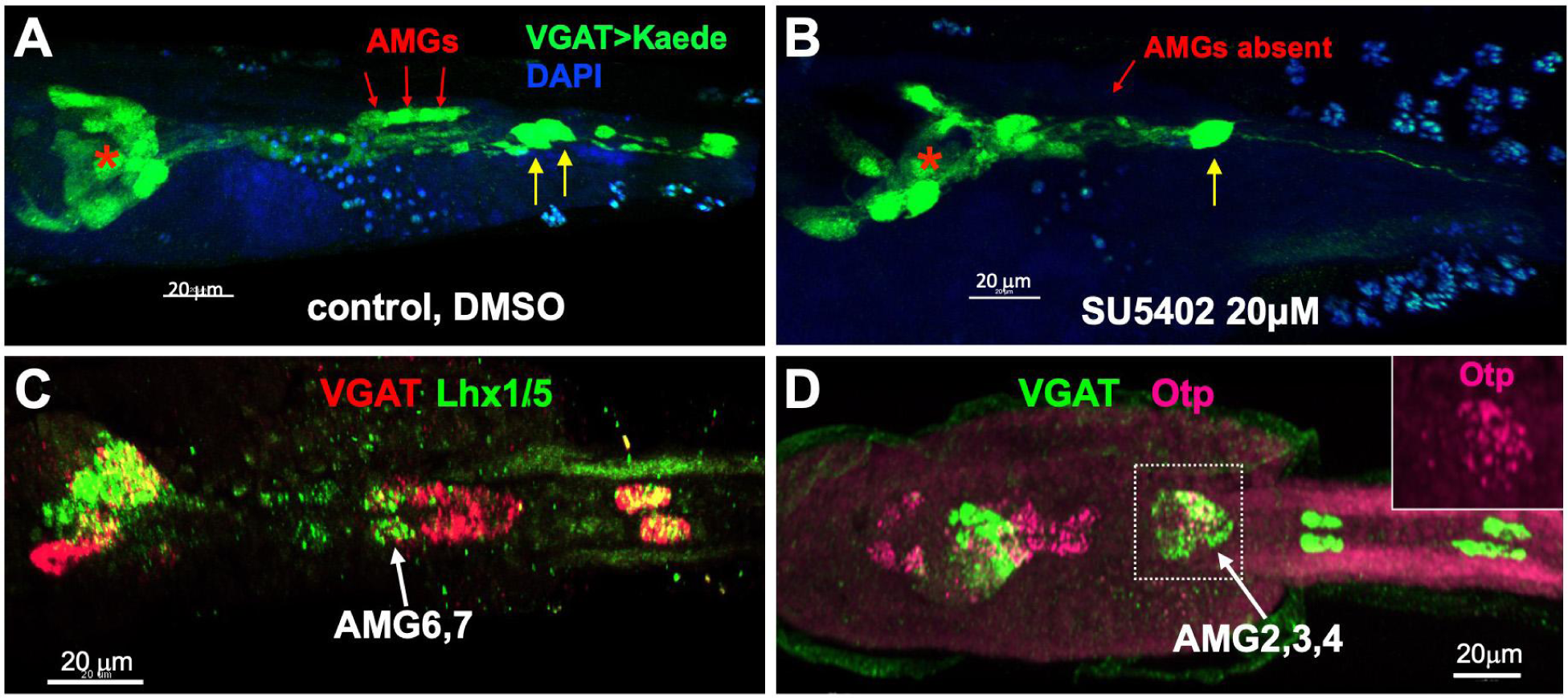
Molecular markers of the AMG group. **A.** Vehicle control for SU5402 experiment (DMSO treatment). The AMGs are labeled by expression of Kaede fluorescent protein driven by the VGAT promoter (red arrows). **B**. SU5402 treatment eliminates the VGAT-expressing AMGs. Red asterisks in panels A and B indicate midbrain VGAT^+^ neurons, while yellow arrows indicate ACINs. **C**. Expression of Lhx1/5 in AMG6 and 7. **D**. Expression of Otp in AMG2, 3 and 4. Both panels C and D show hybridization chain reaction *in situs*.

In addition to the conserved FGF inductive pathway, we also find evidence of conserved gene expression between the cerebellum and the AMG group. However, unlike some cell types and tissues that are defined by the expression of unique transcripts, there are no reports of conserved vertebrate genes that are expressed exclusively in the cerebellum. Nevertheless, there are a number of genes with broad CNS expression that are used as markers for the cerebellum. This includes the genes Lhx1 and Lhx5, which are expressed in Purkinje cells (*36*). We find the *Ciona* ortholog of these two genes, Lhx1/5, is expressed in AMG6 and 7 (Figure 6C). Similarly the gene Otp, which is expressed in the developing mouse cerebellum (*37*), is expressed in AMG2, 3, and 4 [Figure 6D, also see (*17*)]. However, the *Ciona* orthologs of several other broadly-expressed genes that are used as markers of developing vertebrate cerebellum, including *Ptf1a* and *atonal* (*31*), are not expressed in the AMG group (*38*, *39*) (see Discussion).

#### Function and connectomics of the AMG group

A significant difference between the vertebrate cerebellum and the AMG group lies in their respective functions. While the cerebellum functions in the modulation and learning of motor behaviors, the AMG group is a relay and processing center for mechanosensory input. However, vertebrates contain a number of cerebellum-like structures that, like the AMG group, function as mechanosensory relay centers (*40*), suggesting that function of the AMG group may be more similar to that of a cerebellum-like structure. Furthermore, it has been proposed that the cerebellum evolved from a cerebellum-like structure (*41*), providing a plausible evolutionary relationship between the AMG group and cerebellum (see Discussion).

Examination of the circuits of the AMGs, as given by the connectome, suggests a circuit architecture conserved with those in vertebrate cerebellum and cerebellum-like structures, although on a vastly simplified scale, and with some noticeable differences (Figure 7). In *Ciona*, the pATENs project almost exclusively to the single excitatory AMG neuron, AMG5 [Figure 7A and (*30*)]. AMG5 then branches to form synapses with the six inhibitory AMGs. This resembles the relationship between the excitatory granule cells and the inhibitory Purkinje output neurons of the cerebellum, albeit with AMG5 being cholinergic and the granule cells being glutamatergic. Furthermore, in both the AMG group and the cerebellum the output neurons send ascending inhibitory projections, either directly or indirectly, to various regions of the brain. For the AMG group, this includes the prAMG-RNs and eminens neurons of the midbrain (Figure 5C), and the VMG, to which projects are almost exclusively to the MN1 pair. In the cerebellum, the core processing circuit is repeated on a numerically vast scale, while in the AMG group there is only a single circuit consisting of a single excitatory input neuron and six inhibitory output neurons.

**Figure 7.**
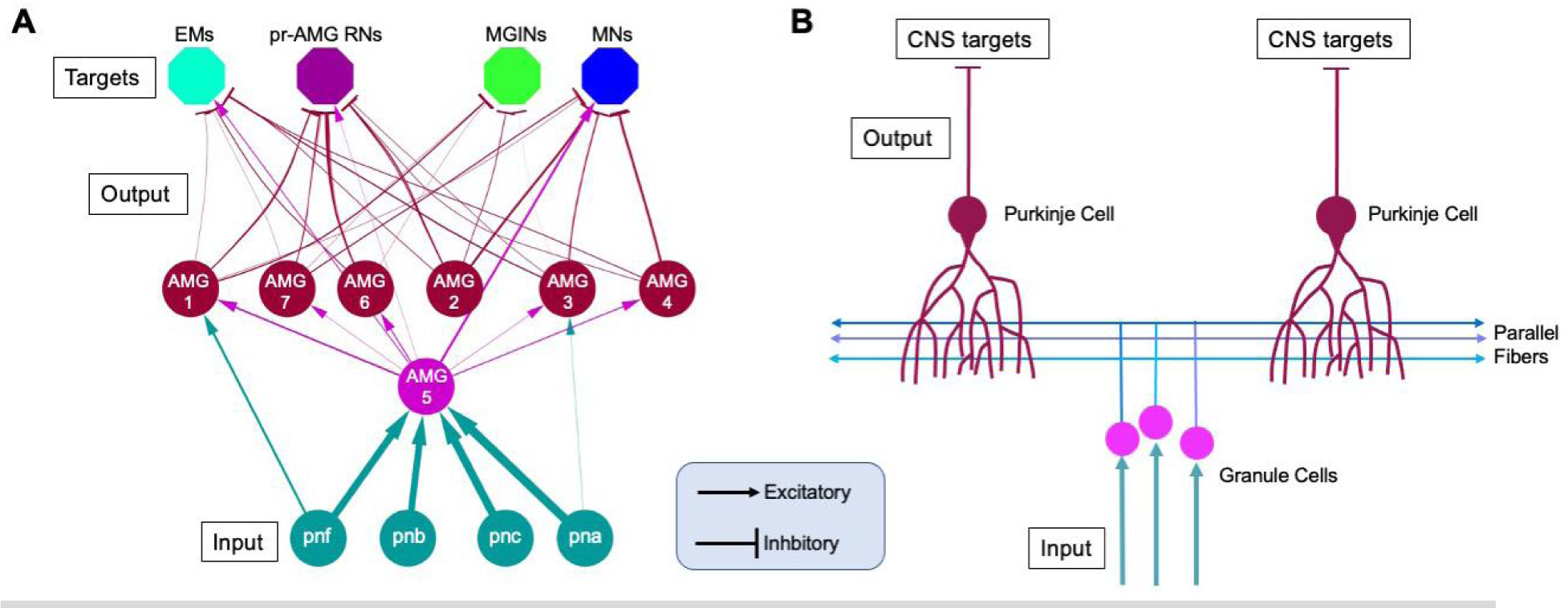
Circuit architecture of the AMG group and cerebellum. **A**. Neural circuit of the AMG group. Input from the mechanosensory pATENs (*pnf*, *pnb*, *pnc* and *pna*) primarily target AMG5 (cholinergic), which then branches to inhibitory output AMG neurons (brown). **B**. Simplified model of the core vertebrate cerebellar cortical circuit. In this circuit, inputs target the excitatory granule cells which then branch (parallel fibers) to the inhibitory output neurons (Purkinje cells).

## DISCUSSION

### The Ciona larval caudal nerve cord and the vertebrate spinal cord

Our description here of motor neurons in the caudal nerve cord supports the homology between the caudal nerve cord and the vertebrate spinal cord. The relatively late development of the caudal nerve cord motor neurons (Figure 1E) may explain why the tail motor neurons had not previously been characterized. In addition to the presence of motor neurons, the highly similar programs of medial intercalation that generate narrow and elongated caudal domains in both tunicate and vertebrate CNS supports their homology (*42*). Additional similarities between the *Ciona* larval caudal nerve cord and the vertebrate spinal cord include the formation of neural crest cells (*43*), and the ventral expression of *sonic hedgehog* along the caudal nerve cord in a pattern resembling that in vertebrate floor plate (*44*). On the other hand, the expression of several genes, such as Hox10 which is expressed in the *Ciona* motor ganglion [Figure 5E and (*45*)] but in the vertebrate spinal cord (*46*), could be taken to argue against homology, it is well documented that expression patterns can be highly divergent between evolutionarily related cell types, particularly in animals as distantly related as tunicates and vertebrates. Thus, while some genes, such as *mnx* appear to have conserved expression and function, the expression of other genes could be misleading with respect to homology. Moreover, as has been discussed (*47*), uncovering deep homologies, such as those between tunicates and vertebrates, will require not only molecular, but also wider morphological and developmental evidence.

### The AMG group, the vertebrate cerebellum, and cerebellum-like structures

The AMG group of neurons makes a distinct domain in the dorsal motor ganglion of *Ciona* larvae. We hypothesize that the similarities between the AMG group and vertebrate cerebellum and cerebellum-like structures are too numerous to be due the convergence of two independently derived structures. These similarities include anatomical location, physical and functional separation from the rest of the motor ganglion/hindbrain, inductive pathway, circuit architecture, and gene expression. The simplicity of the AMG group circuitry differs of course from the enormous complexity of the vertebrate cerebellum or cerebellum-like structure (as does the entire tunicate larval nervous system when compared with those of vertebrates). However, there is also extensive variation in cerebellum complexity among vertebrate classes themselves, with those from mammals being the most elaborate (*48*). Nevertheless, a core circuit of a branching input neuron (AMG5) filling the functional role of the granule cells, and synapsing to inhibitory ascending output neurons (the six VGAT^+^ AMGs) which fill the role of the Purkinje cells, is present. While the VGAT^+^ AMG neurons do not have the extensive arborization that characterizes Purkinje cells (*29*), both Purkinje cells and the VGAT^+^ AMG neurons express Lhx1/5. Moreover, the simplicity of the *Ciona* circuit, with a single input neuron, would argue against the formation of extensive arborizations. One notable difference when comparing the circuits is that AMG5 is cholinergic, while the granule cells are glutamatergic - although both are excitatory. One possibility is that AMG5 and the granule cells have different origins, although it is well documented that homologous neurons can switch neurotransmitters (*49*). It is also worth noting that *Ciona* larvae are devoid of glutamatergic interneurons; rather, the only VGlut^+^ neurons in the larva are sensory (photoreceptors, antennae cells, and peripheral neurons) (*25*). Accordingly, the gene Atoh1, which is expressed in the developing glutamatergic granule cells of the cerebellum (*50*) is expressed in *Ciona* in the pATENs, but not in the AMGs (*39*). However, glutamatergic interneurons are present in cephalochordates (amphioxus), which are basal to both tunicates and vertebrates, suggesting that the lack of glutamatergic interneurons may be a derived feature of tunicates (*51*). Thus, for unknown reasons, tunicates appear to have either lost all glutamatergic interneurons, or changed their neurotransmitter.

The absence of known cerebellum-defining genes means that gene expression may be less informative about homology than other types of evidence. As was introduced above, homologous cell types can have highly divergent expression profiles, even between animals as closely related as the mouse and human (*52*), and evidence of common descent may in some cases be better preserved in morphology and/or function. For example, while the expression patterns of VGAT, Lhx1/5 and Otp are conserved between the cerebellum and AMG group, all are broadly expressed in both *Ciona* and vertebrate CNSs. Another gene product that is used as a marker for vertebrate cerebellum, *zebrin II* [also known as aldolase C; (*53*)], does not appear to have a direct tunicate homolog, rather *Ciona* has a single aldolase gene that is broadly expressed. The transcription factor Ptf1a, which is a key regulator of GABAergic neurons in the cerebellum (*31*), exemplifies the divergence of expression of homologous genes between tunicates and vertebrates. In mammals, Ptf1a, in addition to the cerebellum, is widely expressed, including in the forebrain, pancreas, heart, and spinal cord, and heart (*54*). However, in *Ciona* larvae Ptf1a is expressed exclusively in the *Ciona* coronet cells, which are thought to be related to the vertebrate hypothalamus (*9*).

Functionally, the AMG group, which serves as a relay center for mechanosensory neurons, appears to differ fundamentally from the cerebellum. However, the function of the AMG appears to be like vertebrate cerebellum-like structures. Cerebellum-like structures have the same core circuit architecture as the cerebellum with granule-like and Purkinje-like cells, but overall are simpler (*40*, *55*). Moreover, cerebellum-like structures receive direct input from peripheral electrosensitive and mechanosensitive sensory organs. Examples include a cerebellum-like structure found in the dorsal cochlear nucleus of mammals that receives and processes mechanosensory input from the auditory nerve, and a cerebellum-like structure found in the medial octavolateral nucleus of fish and amphibians that receives input from the mechanosensory lateral line (*56*, *57*). Given the relatively simple structure of cerebellum-like structures, and their widespread distribution, it has been postulated that the cerebellum evolved from a cerebellum-like structure (*41*). This points to a plausible relationship between the cerebellum and the AMG group in which the AMG group reflects an earlier functional role of a cerebellum-like dorsal hindbrain structure. In vertebrates this structure would then have been progressively elaborated and took on new functional roles to become the cerebellum. In that interpretation, in tunicates the same structure may have retained its ancestral role in mechanosensory processing, and may even have become secondarily simplified, as is thought to be a more general characteristic of tunicates (*58*).

A core function of cerebellum-like structures is to act as predictive cancellation filters (*55*, *57*, *59*). Cancellation filters are essential for countering sensory input, known as reafference, that arise from self-stimulation of mechanosensory neurons by the animal’s own motor behavior. The cerebellum-like structures receive proprioceptive afferents that match motor commands which are then used to predict or subtract the self-generated signals (Figure 8A). The predictive cancelation inputs consist of both excitatory afferents that convey both the external stimulus and the self-generated noise, and inhibitory interneurons that are responsive only to the self-generated noise. Examination of the *Ciona* connectome reveals a plausible predictive cancelation filter functionality in the AMG group (Figure 8B). In addition to receiving direct mechanosensory input from the pATENs at AMG5, the AMGs are postsynaptic to the four *bipolar tail neurons* (BTNs), which function as relay interneurons for the mechanosensory *dorsal caudal epidermal neurons* (DCENs) (Figure 8C). The BTNs have mixed valence, with the anterior two (BTN1 and 2) being VGAT^+^, while the posterior two (BTN3 and 4) are VACHT^+^ (Figure 8D). The output from the BTNs, which should reflect the motor commands to the tail, could provide the necessary information for a predictive cancellation filter. While a predictive cancellation filter in the *Ciona* AMGs is conjectural, it is likely that *Ciona* mechanoreceptors would face the same self-stimulation problem as in other animals. Moreover, the circuitry of the AMG group, which includes direct excitatory sensory input from the pATENs which funnel through AMG5 to the inhibitory output AMGs, as well as indirect excitatory and inhibitory input from the DCENs via the BTNs, appears to be unnecessarily complicated if the AMG group were simply acting as a relay center for mechanosensory input, rather than as a processing center.

**Figure 8.**
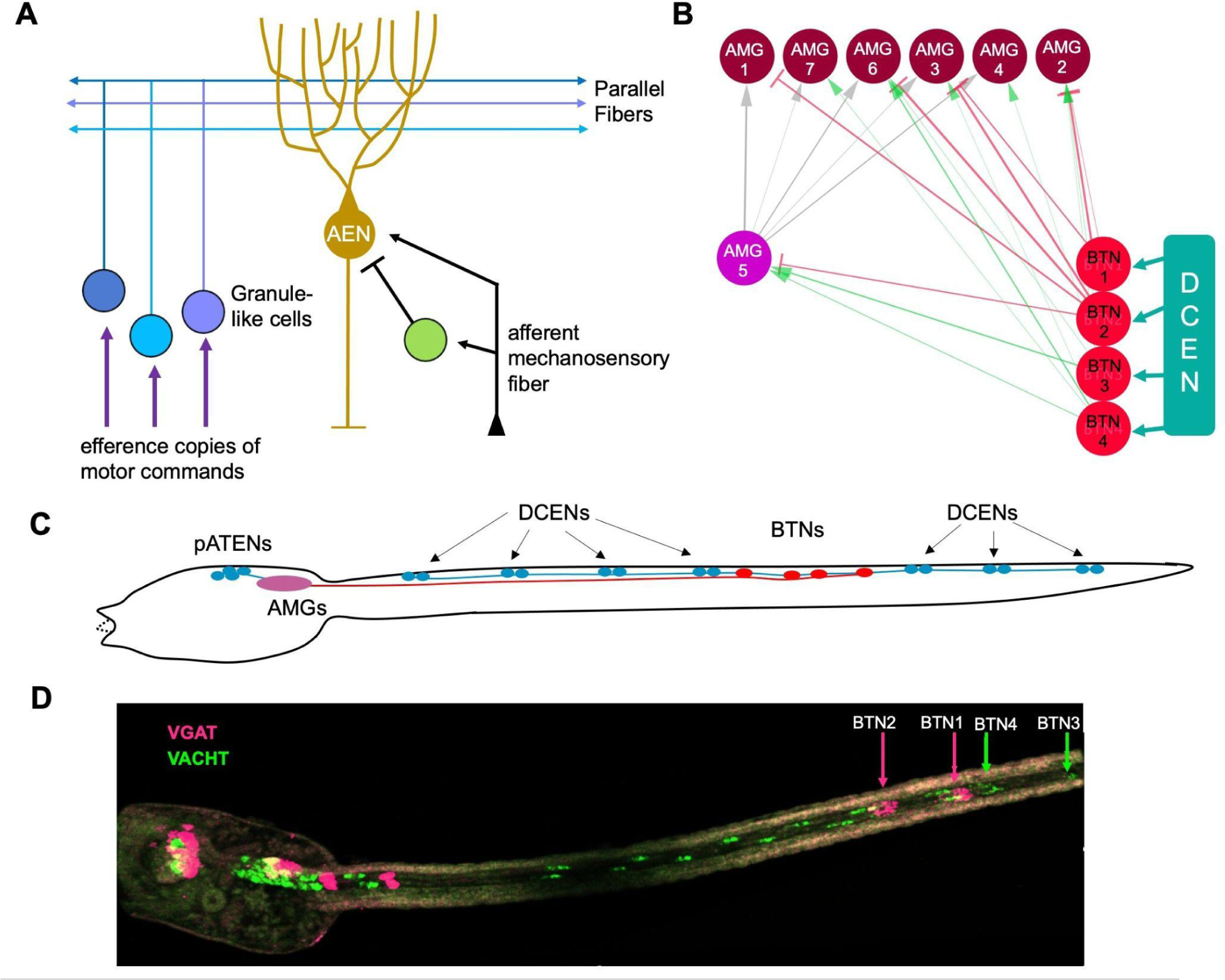
Predictive cancelation circuits. **A**. Simplified depiction of a predictive cancelation circuit in the dorsal octavolateralis nucleus [based on (*55*)]. **B**. Synaptic input from the BTNs to the AMG group. **C**. BTNs labeled by in situ hybridization for VACHT (green) and VGAT (magenta). **D**. Relative positions of the pATENs, DCENs, AMGs and BTNs in a *Ciona* larva. Abbreviations: AEN = Purkinje-like *ascending efferent neuron*; AMG = *ascending motor ganglion interneuron*; pATEN = *posterior apical trunk epidermal neuron*; DCEN = *dorsal caudal epidermal neuron*; BTN = *bipolar tail neuron*.

### Summary

The compiled evidence on the development, morphology, neuron types and function of the *Ciona* caudal nerve cord are consistent with this and the vertebrate spinal cord originating from a common structure with spinal cord-like features. The third group of chordates, the cephalochordates, which are basal to both tunicates and vertebrates, likewise has a spinal cord (*60*), indicating that this structure was present at the base of the chordates. As we have done in a previous publication, we therefore propose that the tunicate caudal nerve cord simply be called the spinal cord. By contrast, we do not think it would be correct to call the AMG group a cerebellum. Rather, we speculate that the AMG group derives from an ancestral cerebellum-like structure that preceded the vertebrate cerebellum. Whether this structure was also found in the common ancestor of all chordates is not clear. No cerebellum or cerebellum-like structure has been described in cephalochordates. However, the properties of the AMG group that distinguish it from the rest of the motor ganglion, and that point to cerebellum-like homology, were not evident until the connectome dataset was available. Similar information is currently lacking for any cephalochordata species. The presence of a cerebellum-like structure in the motor ganglion, along with the presence of a clear spinal cord homolog, lead us to propose that this structure can confidently be called a hindbrain homolog, rather than a homolog of both the hindbrain and spinal cord. Finally, these newly recognized homologies between tunicate and vertebrate CNSs reinforces the utility and validity of tunicates as a model, aided by their simplicity, described connectome, and extensive behavioral repertoire, for exploring neural mechanisms that are shared by all chordates.

## MATERIALS AND METHODS

### In situ hybridization

To label mRNA fluorescently in whole mount *Ciona* larvae or embryos, the hybridization chain reaction (HCR) in situ method (Molecular Instruments) was followed, as previously described (*25*). Genes for which probe sets were designed and their sequence identifiers are as follows: vesicular acetylcholine transporter (VACHT; NM_001032789.1), motor neuron homeobox (MNX; KH2012:KH.L128.12) lim-domain homeobox 1/5 (LHX/5; KH2012.L107.7), vesicular GABA transporter (VGAT; NM_001032573.1), and Orthopedia (OTP; XM_009862516.3).

#### Chrimson

A plasmid encoding *mCherry tagged-chrimson* channelrhodopsin (*61*) driven by the *Ciona* VACHT *cis*-regulatory region (*19*) was constructed by amplifying the promoter region of VACHT from pSPCiVACHTC (obtained from CITRES, marinebio.nbrp.jp/ciona/index.jsp) and inserting this into TRE::ChrimsonR:mCherry (obtained from Addgene, clone #99207) at the Sac1 and EcoR1 sites. The primers used to amplify the VACHT promoter are AGagctcTTAAGCACACGTTCAAAAAT and TCGAATTcGATGAACAATAAAGTAG.

One-cell stage embryos were electroporated with 45 μg of plasmid as previously described (*22*). Control embryos received no electroporation. At early tailbud stage embryos were bisected into head and tail regions using the tip of a 30-gauge hypodermic needle as a knife. At the larval stage, the bisected tails were placed into the wells of depression slides with filtered seawater containing 0.05% methylcellulose. Time-lapse videos were recorded at 8.9 frames/second with a Hamamatsu ORCA-ER camera fitted with a Navitar 7000 zoom lens and a Tiffen #47 blue filter. Imaging sessions were for 30 seconds with continuous 0.6 mW/cm^2^ illumination from a 470 nm LED lamp (ThorLabs). At the midpoint of the imaging session (*i.e.*, at 15 seconds) a 590 nm LED lamp (Thorlabs) set to 5.3 mW/cm^2^ was turned on. Time-lapse videos were analyzed using ImageJ. Following time-lapse capture, tails were immunolabeled with anti-mCherry antibodies and imaged.

#### Immunolabeling

Labeling with antibodies was performed as previously described (*12*). Primary antibodies used were as follows: mCherry (rat; life Technologies), Kaede (rabbit; MBL), and GFP (mouse; Invitrogen). Appropriate fluorescent Alexa Fluor secondary antibodies were purchased from Invitrogen.

#### SU5402 treatment

SU5402 (Sigma) was diluted in DMSO to make a stock solution at 10µg/µL, which was diluted to a working solution ranging from 20-40µM. DMSO-only served as a negative control. The drug was added to seawater-filled dishes containing late-neurula stage embryos which resulted from a cross between wild type eggs and stable transgenic VGAT>Kaede sperm. Thirty minutes after addition of the drug, the embryos were washed four times with filtered seawater, then allowed to reach the larval stage, when they were fixed for immunolabeling.

### *Ciona* Connectome

Analysis of *Ciona* connectomic data, and generation of connectivity diagrams, was performed using Cytoscape (*62*). The connectivity matrix is available for download from (*5*). Reconstruction of *Ciona* CNS domains from the connectome project serial-section EM images were generated using the Cionectome Viewer on Morphonet (https://morphonet.org/).

## Acknowledgments

We thank Yishen Miao for bioinformatics help.

## Supplemental Material

**Video 1**. Larval tails expressing VACHT>Chrimson-mCherry (and non-expressing controls tails). Midway through video the samples are illuminated with a 590 nm LED lamp. Movie plays at 5-times normal speed.

**SFigure 1.**
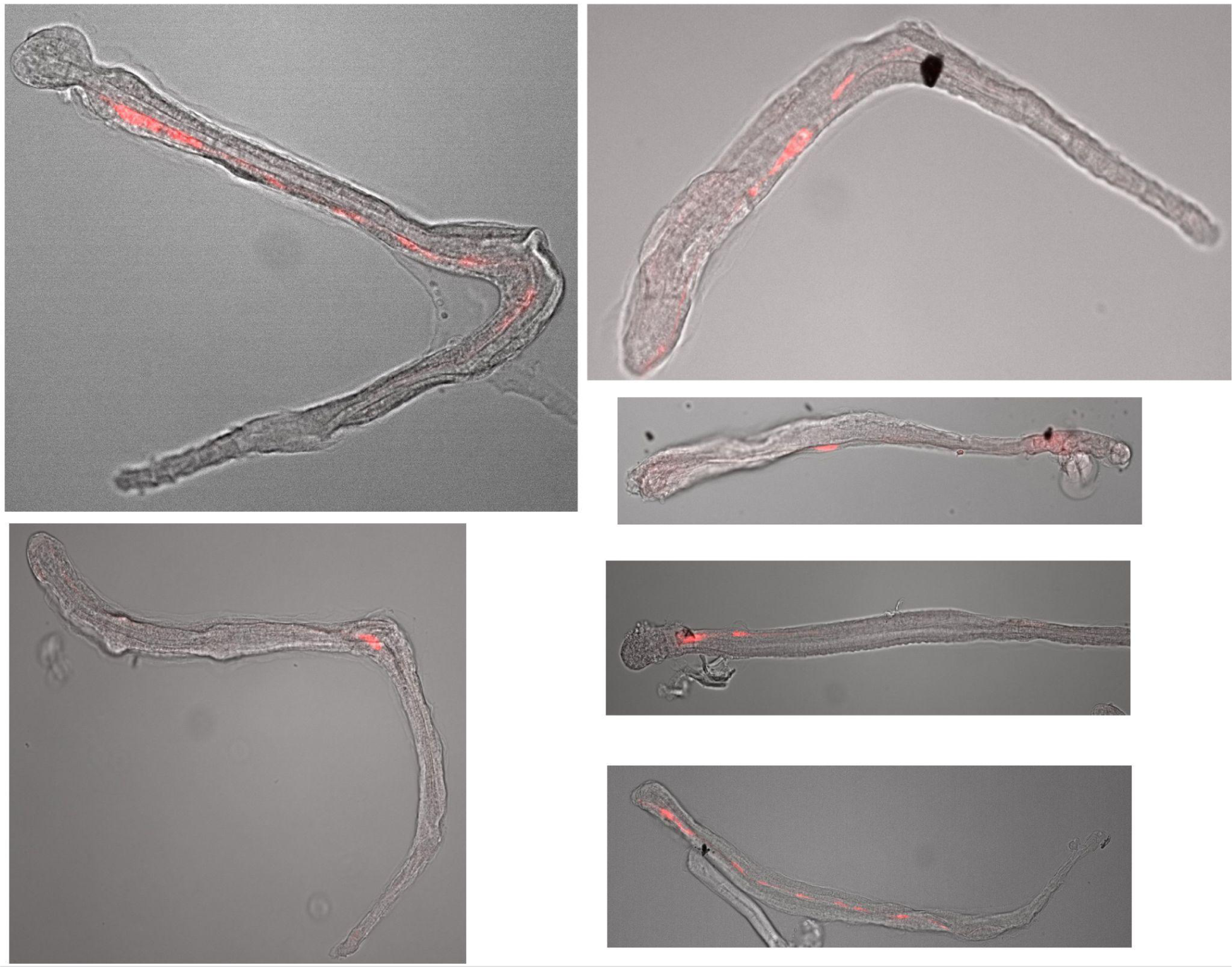
Immunolabeled *Ciona* larval-stage tail fragments expressing VACHT>Chrimson-mCherry. The samples were immunolabeled with a rat anti-mCherry primary antibody and an Alexa Fluor 546 goat anti-rat secondary.

**SFigure 2.**
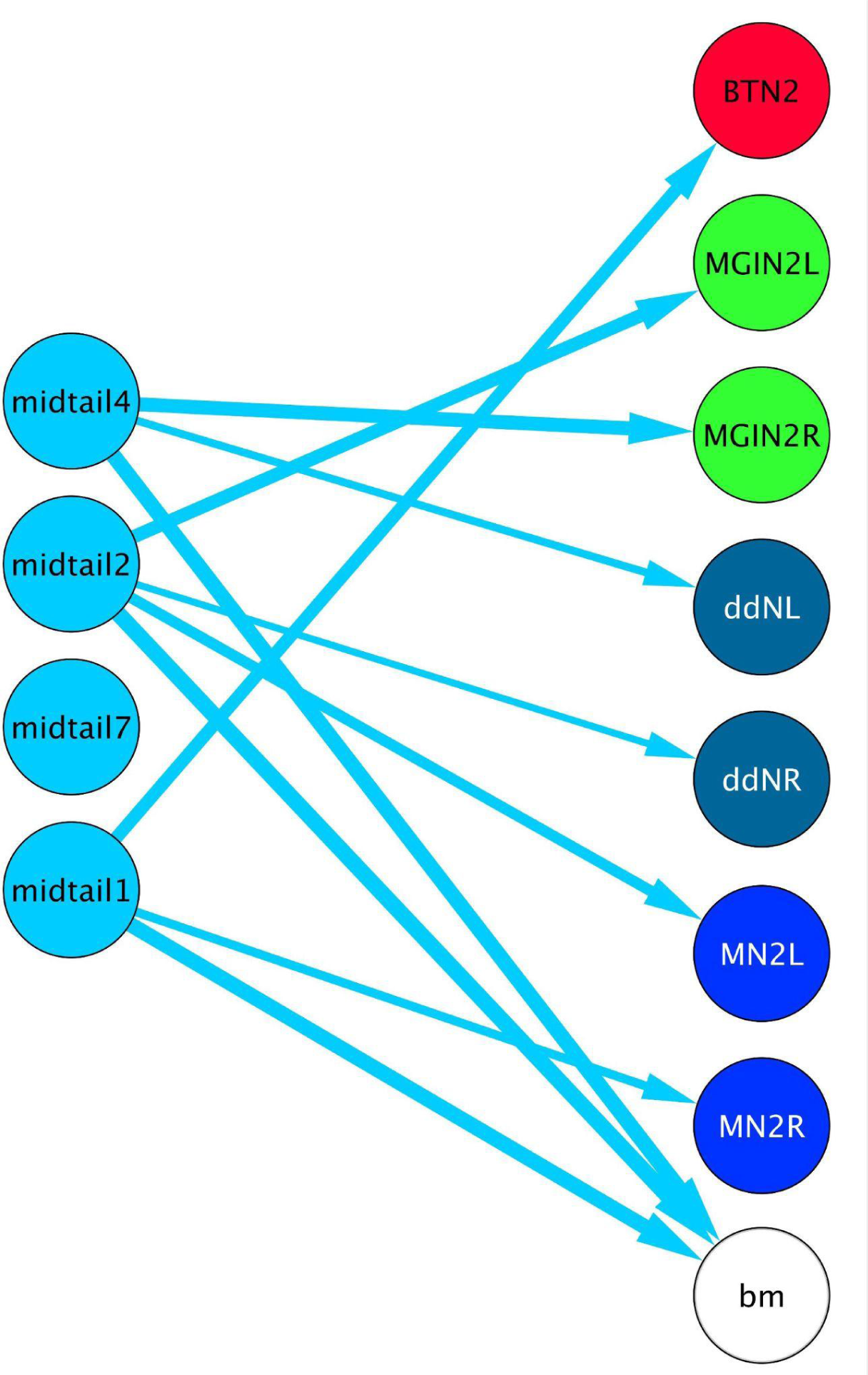
Synaptic targets of the midtail neurons (excluding muscle cells), as given by the *Ciona* connectome.

