## Supplemental Figures for "Deep homologies in chordate caudal central nervous systems"

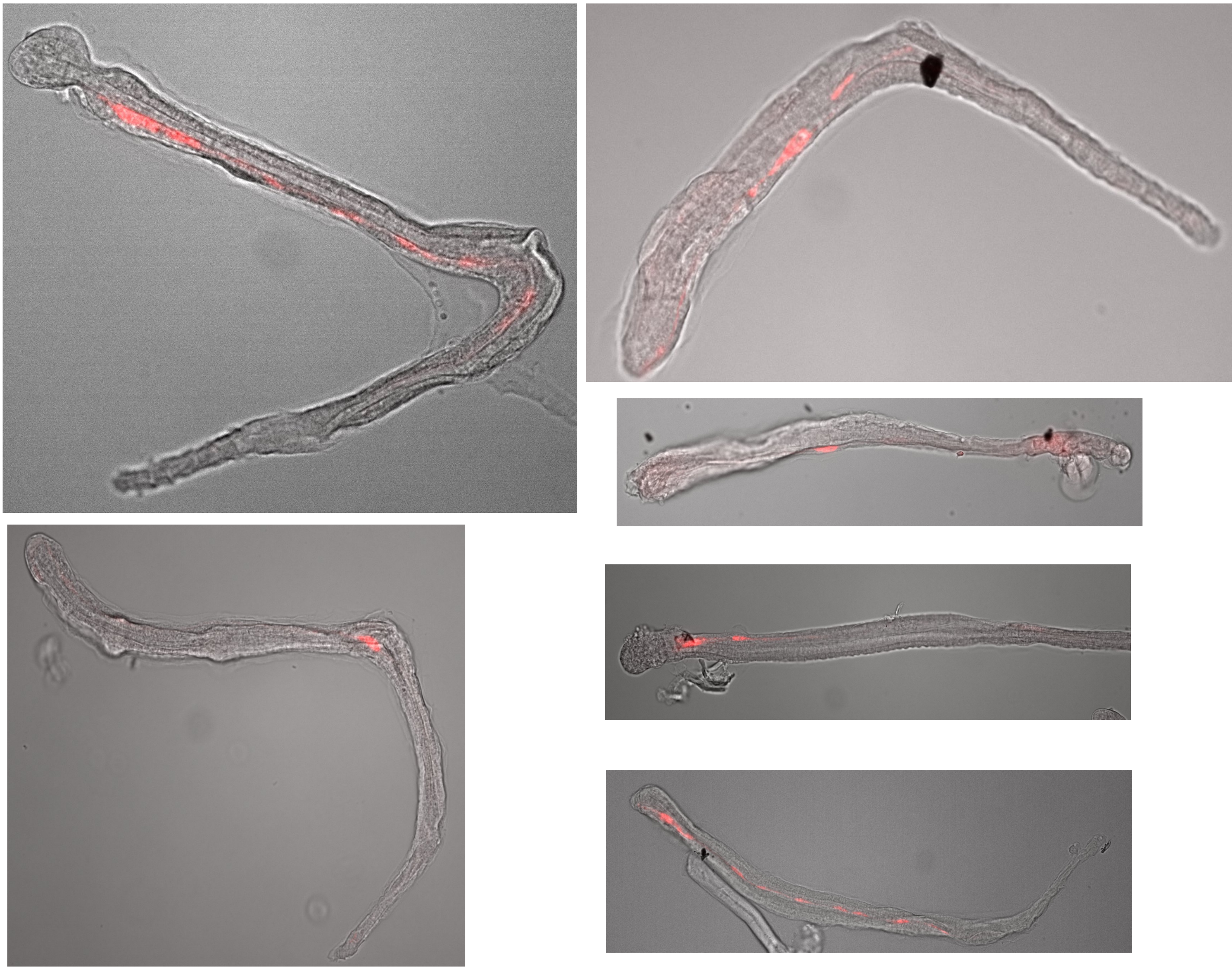

**SFigure 1**. Immunolableing of *Ciona* larval-stage tail fragments expressing VACHT>Chrimson-mCherry. The samples were immunolabled with a rat anti-mCherry primary antibody and an Alexa Fluor 546 goat anti-rat secondary.

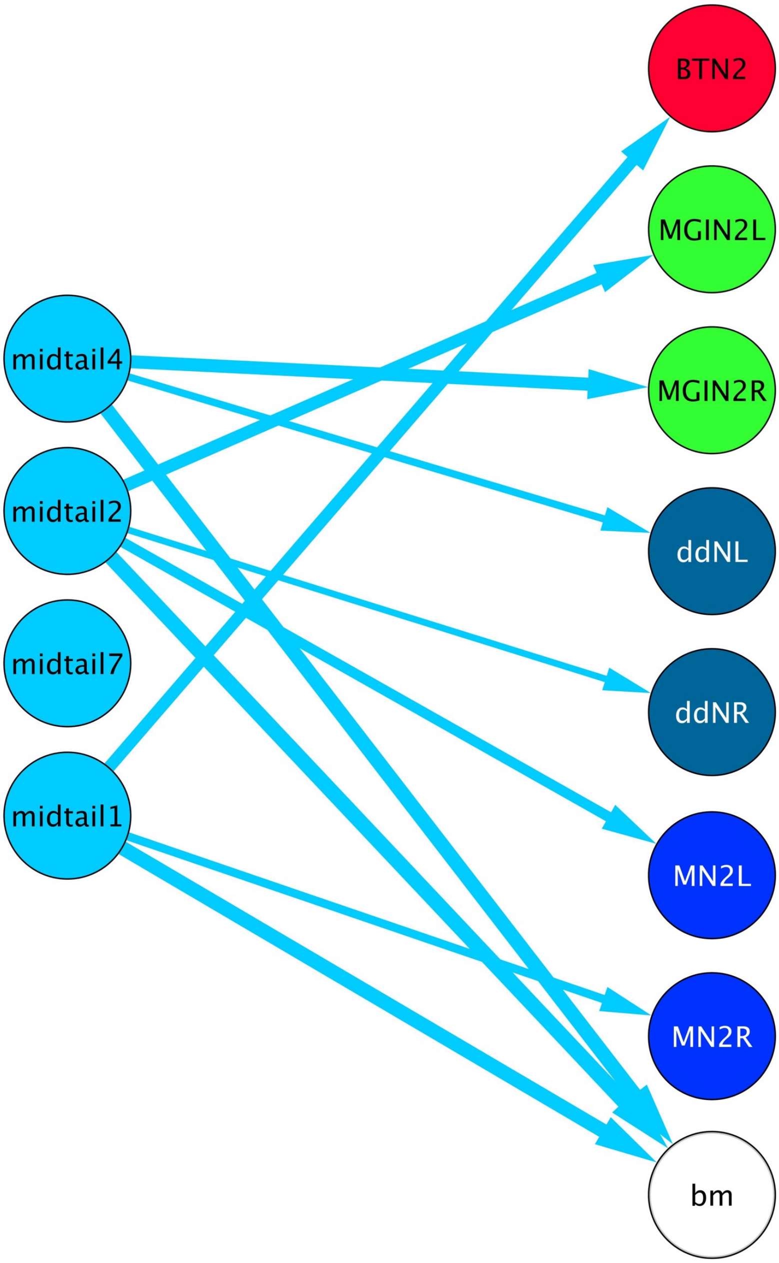


**SFigure 2**. Synaptic targets of the midtail neurons (excluding muscle cells), as given by the *Ciona* connectome
